# Genetic architecture of transmission stage production and virulence in schistosome parasites

**DOI:** 10.1101/2020.12.23.424224

**Authors:** Winka LE CLEC’H, Frédéric D. Chevalier, Marina McDew-White, Vinay Menon, Grace-Ann Arya, Timothy J.C. Anderson

## Abstract

Both theory and experimental data from pathogens suggest that the production of transmission stages should be strongly associated with virulence, but the genetic bases of parasite transmission/virulence traits are poorly understood. The blood fluke *Schistosoma mansoni* shows extensive variation in numbers of cercariae larvae shed and in their virulence to infected snail hosts, consistent with expected trade-offs between parasite transmission and virulence. We crossed schistosomes from two populations that differ 8-fold in cercarial shedding and in their virulence to *Biomphalaria glabrata* snail hosts, and determined four-week cercarial shedding profiles in F0, F1 and 376 F2 progeny from two independent crosses in inbred snails. Sequencing and linkage analysis revealed that cercarial production is polygenic and controlled by five QTLs. These QTLs act additively, explaining 28.56% of the phenotypic variation. These results demonstrate that the genetic architecture of key traits relevant to schistosome ecology can be dissected using classical linkage mapping approaches.

## INTRODUCTION

Parasites face a central problem: how to maximize transmission to the next host. This has driven the evolution of a wide variety of lifecycle features to facilitate parasite transmission (Criscione et al., 2020; Vasudevan et al., 2015). However, perhaps the most common transmission strategy is to produce vast numbers of infective stages (Loker and Hofkin, 2015, p. 20). This brute force approach exploits host resources for parasite growth and reproduction, while infective stages may also cause damage as they exit the host to reach the environment (Jensen et al., 2006, p. 200; Le Clecʼh et al., 2019). Hence, there is a strong expectation that transmission stage production will result in collateral damage to the host, but also that high levels of host mortality will constrain evolution of very high levels of parasite virulence. There is a large body of theoretical work examining this relationship between transmission stage production and virulence evolution (Alizon et al., 2009; Anderson R.M., 1982; Frank, 1996) and empirical studies provide compelling, but not universal support, for this model (Acevedo et al., 2019; Alizon and Michalakis, 2015). However, there is limited understanding of the genes and genetic architecture underlying transmission stage production and virulence on which selection can act.

Schistosome parasites provide a useful system for examining the genetic basis of transmission/virulence related traits. These parasitic organisms are well suited for genetic studies because (i) the complete lifecycle can be maintained in the laboratory using rodent definitive hosts and freshwater snail intermediate hosts, (ii) parasites have separate sexes which simplifies staging of efficient genetic crosses in the laboratory, (iii) thousands of progeny are produced which provides good statistical power, and (iv) experimental work over the last 75 years has revealed heritable genetic variation in multiple biomedically important traits (Anderson et al., 2018) such as drug resistance (Cioli et al., 1992; Greenberg, 2013; Melman et al., 2009; Mwangi et al., 2014; Valentim et al., 2013), chronobiology (Théron, 2015; Théron and Combes, 1983, 1988), host specificity (Files and Cram, 1949; Kalbe et al., 2004; Mitta et al., 2017; Rollinson et al., 2001; Theron et al., 2014), and virulence (Davies et al., 2001; Gower and Webster, 2004; Webster et al., 2004). Furthermore, the *Schistosoma mansoni* genome is fully sequenced and assembled (Berriman et al., 2009; Protasio et al., 2012), and a growing molecular toolkit including molecular sexing tools (Chevalier et al., 2016; Gasser et al., 1991), RNAi (Krautz-Peterson et al., 2010), transfection (Mann et al., 2014; Rinaldi et al., 2012), CRISPR (Ittiprasert et al., 2019; Sankaranarayanan et al., 2020; You et al., 2021) and a suite of cell biology tools (Collins and Collins, 2017; Wang et al., 2020; Wendt et al., 2020; Wendt and Collins, 2016) improves our ability to link phenotype with genotype. Furthermore, we can also control the genetics of the snail host by generating inbred snail lines. *B. glabrata* is hermaphrodite, so inbred snail lines can be developed by isolating individual snails and serial inbreeding (Bonner et al., 2012).

In addition of being a tractable model for genetic studies, schistosomes are also devastating human parasites. Three major schistosome parasite species infect over 200 million people in 78 countries (Verjee, 2019; World Health Organization, 2019). These parasites have a complex lifecycle, involving a freshwater snail (intermediate host) and a mammal (definitive host). When parasite eggs are expelled with mammal feces or urine in freshwater, miracidia larvae hatch and actively search for its snail vector. Larvae penetrate the snail head-foot, differentiate into sporocysts that then asexually proliferate to generate daughter sporocysts. These actively consume snail tissue (hepatopancreas and ovotestis), castrating the host snail, and parasite sporocysts can comprise up to 60% of the tissue within infected snails (Le Clecʼh et al., 2019). The daughter sporocysts release cercariae, the mammal-infective larval stage of the parasite. Hundreds to thousands of these motile cercariae exit through the snail body wall and are released into freshwater. Exit through the body wall results in leakage of hemolymph and damage to the snail. This quantitative transmission and virulence related trait – numbers of cercarial produced from the intermediate snail host – is the focus of this paper.

There is strong evidence from laboratory selection experiments that this cercarial shedding is heritable: in just three generations parasites selected for low of high numbers of cercariae showed rapid divergence in this phenotype (Davies et al., 2001; Gower and Webster, 2004). We have described distinctive life-history strategies in laboratory schistosome populations. In two parasite populations originating from Brazil we observed: i) a “boom-bust” strategy characterized by high transmission measured in term of cercarial production (the number of larvae released from the intermediate snail host) and high virulence to the snail intermediate host resulting in short duration infections, compared with ii) a “slow and steady” strategy characterized by low transmission (few cercariae released), resulting in low virulence to the snail host and a long duration of infection (Le Clecʼh et al., 2019). The high shedding genotype develops larger sporocyst stages than the low shedding genotype, although this is insufficient to explain the difference in cercarial shedding from these two parasites. These patterns are observed when using the same inbred snail line, ruling out an impact of host genetics. Our central goal was to decipher the genetic architecture of a transmission/virulence-related trait in a biomedically important helminth parasite, to determine whether cercarial shedding is under monogenic or polygenic control, and ultimately to understand the cellular pathways on which selection acts to modulate this trait.

Linkage mapping has previously been used to examine a monogenic drug resistance trait (oxamniquine resistance) in *S. mansoni* (Chevalier et al., 2014; Valentim et al., 2013). Here we extend this approach to examine a life-history trait: we conducted reciprocal genetic crosses between *S. mansoni* individuals from two laboratory parasite populations originating from Brazil: the SmBRE (Low Shedder - L) population and the SmLE (High Shedder - H) population (Figure 1). Using classical linkage mapping, we showed that cercariae production in *S. mansoni* parasite is polygenic, with additive variation at 5 different QTLs explaining 28.56% of the variation in cercarial production.

**Figure 1:**
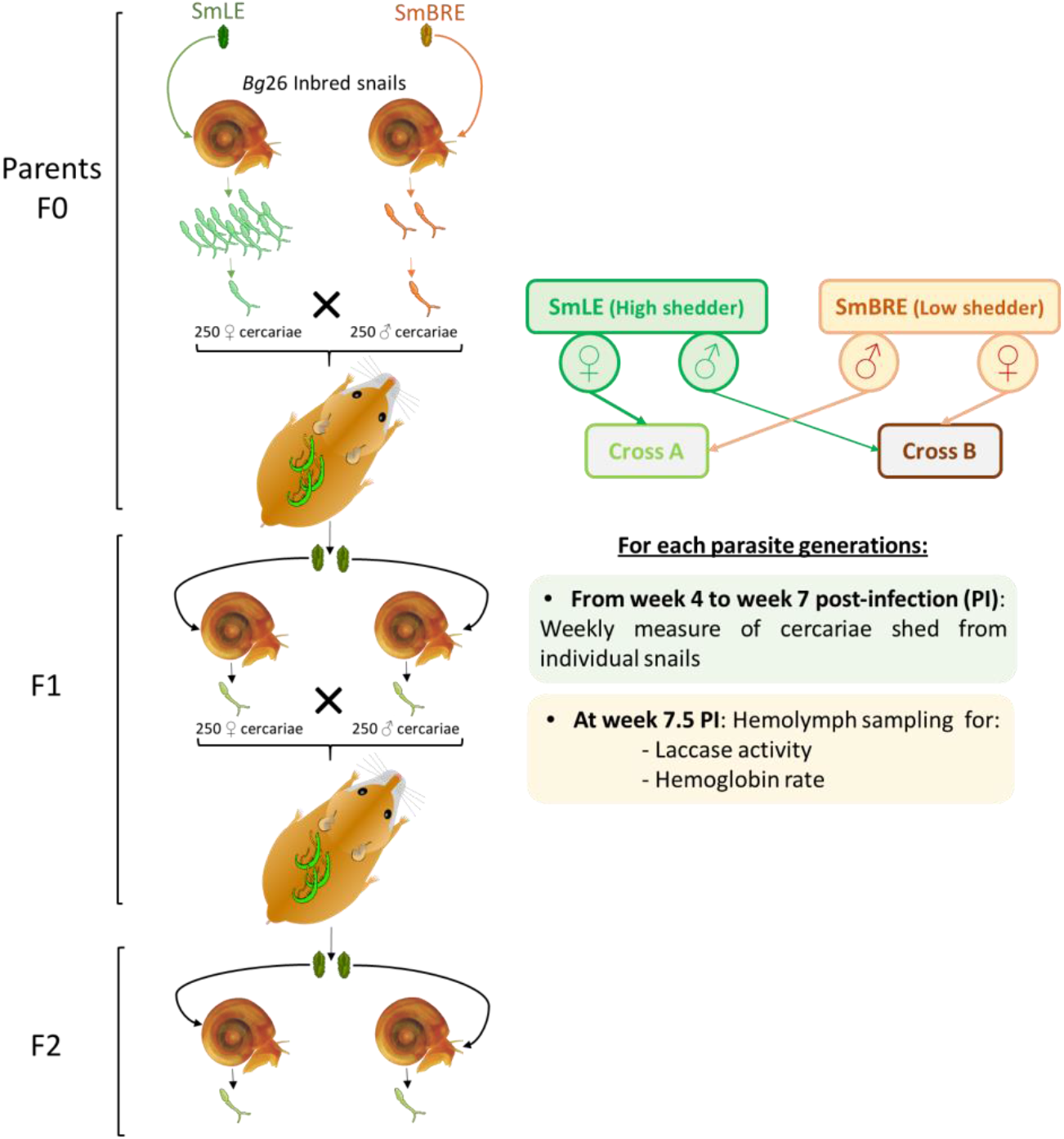
Outline of the genetic cross experimental design. We performed two independent reciprocal genetic crosses (i.e crosses A and B, to account for potential sex specific effect(s)) between single genotypes of SmLE-H and SmBRE-L *Schistosoma mansoni* parasite populations. These two populations’ exhibit striking differences in term of transmission stage production (i.e. number of cercariae produced). For each parasite generation (i.e F0 parental populations, F1s and F2s progeny), we exposed individual *Biomphalaria glabrata* Bg26 inbred snail to single miracidium from either the SmLE-H or SmBRE-L populations for F0 (*n*=192/population), or the F1 progeny (*n*=288/cross), or the F2 progeny (*n*=1000/cross). For each infected snail and each generation of *S. mansoni* parasites, we measured transmission stage production during 4 weeks of the patent period (week 4 to 7 post-infection). We also evaluated the virulence of these generations of crossed parasites by measuring the daily snail survival during the patent period. After 7.5 weeks post-infection, surviving infected snails were bled and we measured the total laccase-like activity as well as the hemoglobin rate in the collected hemolymph samples.

## RESULTS

### *Variation in transmission stage production in* S. mansoni *crosses*

We conducted two three-generation (F0 to F2) genetic crosses between high (H) shedding parasite genotypes SmLE-H and low (L) shedding genotypes SmBRE-L. We paired a male SmBRE x a female SmLE (cross A) and a female SmBRE x a male SmLE (cross B) to allow examination of sex specific inheritance. The two *S. mansoni* populations, SmLE-H and SmBRE-L, both originated from Brazil, and show striking difference in cercarial production (Le Clecʼh et al., 2019). We used the same inbred *B. glabrata* population (Bg26) in these experiments to minimize the impact of the host genetics, because cercarial shedding can be influenced by snail genotype (Jones-Nelson et al., 2011). Furthermore, we infected snails with single parasite miracidia, so quantitative measure of cercarial production by infected snails can be related to a single parasite genotype.

Using weekly measures of cercarial shedding over 4 weeks, we observed that SmLE (H) population shed 8-fold more cercariae than SmBRE (L) (mean (±se) cercariae per shedding: SmBRE (L): 284±19 vs SmLE (H) 2352±113; Wilcoxon test, *p* < 2.2 × 10^™16^, Figure 2A-B F0; Le Clecʼh et al., 2019). However, the infectivity of these two parasite population in Bg26 snails was identical (i.e. 26% of the snails exposed with one miracidia from either SmLE-H or SmBRE-L were infected).

**Figure 2:**
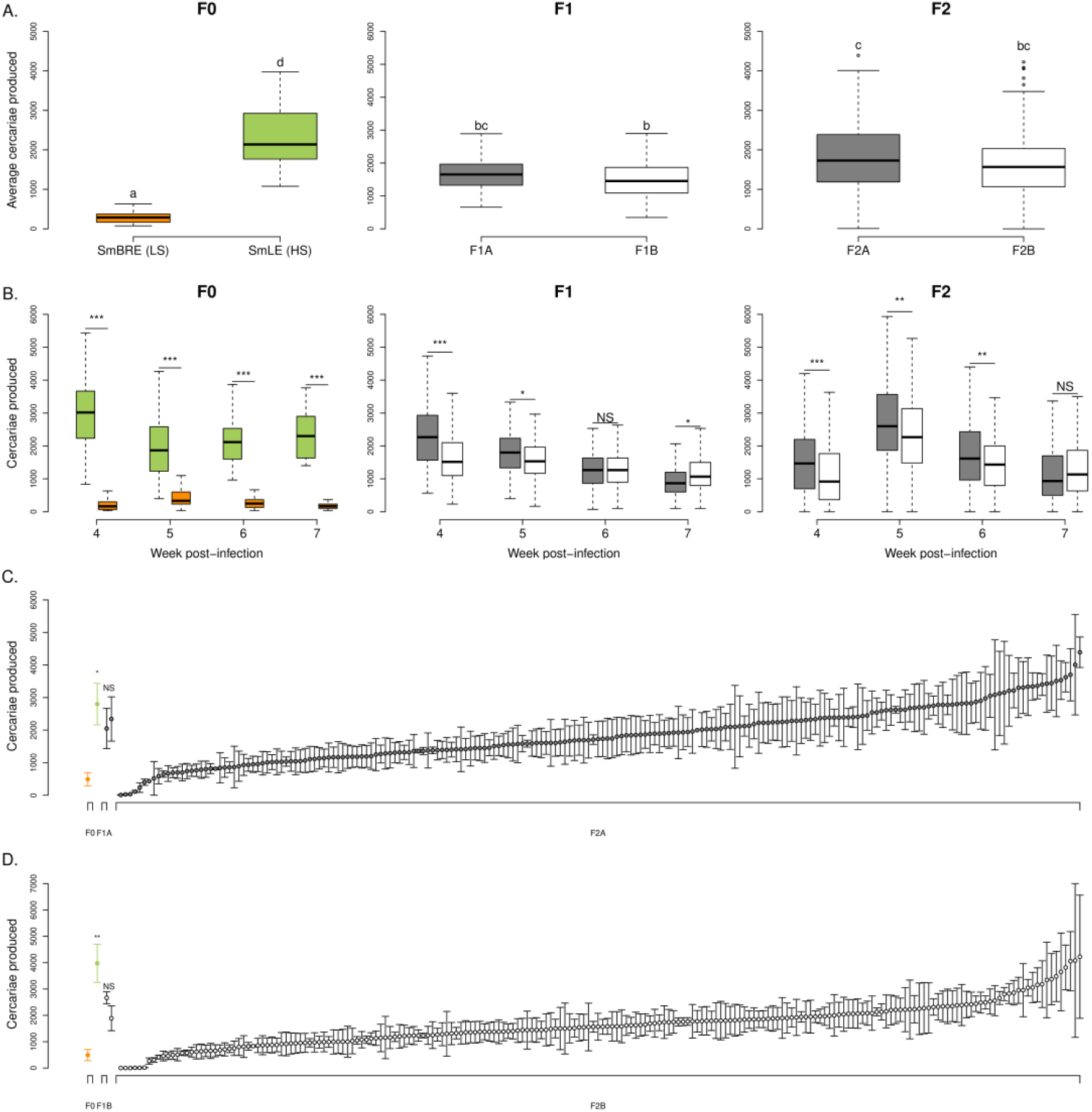
Transmission stage production of two *Schistosoma mansoni* parental populations (i.e. SmLE-H and SmBRE-L) and their progeny (i.e. F1s and F2s). **(A)** Difference in the average of cercariae produced by SmLE-H and SmBRE-L *S. mansoni* populations during 4 weeks of the patent period (week 4 to 7 post infection). SmLE-H population is shedding more cercariae than SmBRE-L population (Le Clecʼh et al., 2019). For both crosses A and B, F1s populations exhibit intermediate phenotype in terms of cercarial production, compared to F0, while the average production of cercariae by F2s encompass both parental and F1s distributions. Parasite populations or generations not connected by the same letter are significantly different (post-hoc test). **(B)** Difference in the number of cercariae produced by *S. mansoni* parental populations (i.e. SmLE-H and SmBRE-L) and progeny (i.e. F1 and F2) measured by week (week 4 to 7 post infection). **(C)** Distribution of the cercarial production (mean ± SE) over the 4 weeks of the patent period for the parents (F0) of the cross A, the F1 parents (F1A) and all the 204 F2A progeny by rank order. **(D)** Similar distribution plot of the cercarial production for the parents (F0), the F1 parents (F1B) and all the 204 F2B progeny for the cross B by rank order. For both crosses **(C-D)**, L and H F0 parents exhibited striking differences in cercarial production (cross A: Welsh t-test, *p*= 0.031; cross B: Welsh t-test, *p*= 0.013) while F1 showed an intermediate phenotype and F2 encompassed both parental distribution with a gradation from L to H phenotype. NS: No significant difference in cercarial production between the two considered groups, * *p* < 0.05, ** *p* ≤ 0.02, *** *p* ≤ 0.002.

We measured the cercarial production of the two *S. mansoni* F1 populations (i.e. F1A for cross A and F1B for cross B). For both crosses, F1 parasite populations shed intermediate numbers of cercariae compared to F0 (i.e. H and L parental populations) (Kruskal-Wallis test followed by pairwise Wilcoxon Mann-Whitney post-hoc test, *p* < 2.2 × 10^™16^) and showed limited variation in shedding number (Figure 2A). Schistosome gender impacted the number of cercariae produced by the F1A parasite population with male genotypes producing significantly less cercariae than female ones (one-way ANOVA, *p*= 0.0038, Supplementary figure 1A). We did not see any effect of parasite gender in the F1B progeny (Supplementary figure 1B). Moreover, F1 parasites from cross A produced significantly more cercariae than those from cross B (Wilcoxon test, *p* = 0.0048, Figure 2A-B).

The F2 parasite progeny from both crosses showed extensive variation in numbers of cercariae shed that encompassed the range seen in the two parental distributions (Kruskal-Wallis test followed by pairwise Wilcoxon Mann-Whitney post-hoc test between F0 populations (i.e. SmLE-H and SmBRE-L) and F2s (i.e. F2A and F2B), *p* < 2.2 × 10^™16^, Figure 2A F2). In F2s from both crosses, we observed parasites that produced very few cercariae, like the L parent, some that produced cercariae at an intermediate level, like F1s, and some that produced large numbers of cercariae like the H parent (Figure 2C and 2D). F2 progeny from cross A shed more larvae than cross B (Wilcoxon test, *p* = 0.0089, Figure 2A-B F2). The F2 progeny have an average cercarial shedding profile across weeks that mimic the L parental profile with a peak in cercarial production arising on the second shedding. However, the average intensity of cercarial production is closer to the H parent (Figure 2A-B). We observed a significant impact of schistosome gender in cross B (Kruskal-Wallis test, *p*=0.041, Supplementary figure 1C) but not in cross A (Kruskal-Wallis test, *p*=0.719, Supplementary figure 1D) for the F2s with male genotypes producing significantly less cercariae than female ones.

### *Linkage analysis of transmission stage production in* S. mansoni *parasites*

We sequenced the whole genome (363 Mb) from parents and the exome (15 Mb) of F1s and F2s progeny from each cross (Figure 1). We found 10,543 (cross A) and 8,779 (cross B) SNPs fixed for alternative alleles (i.e. fully informative markers) in the exome. A classical linkage analysis using as phenotype the average quantities of cercariae per snail (i.e. schistosome genotype) revealed three major quantitative trait loci (QTL) involved in transmission stage production: on chr. 1 (LOD= 5.63), chr. 3 (LOD= 8.16) and chr. 5 (LOD= 6.37) (Figure 3A). This finding is consistent with the minimum number of loci calculated using the Castle-Wright Estimator (Castle, 1921; Cockerham, 1986; Hedrick, 1983; Lande, 1981; Wright and Morton, 1968), which estimates the minimum number of loci determining trait variation from patterns of segregating phenotypic variation. This estimated a minimum of 3.03 loci involved for cross A and 1.85 for cross B. Gender taken as a covariate did not impact the result of the linkage analysis.

**Figure 3.**
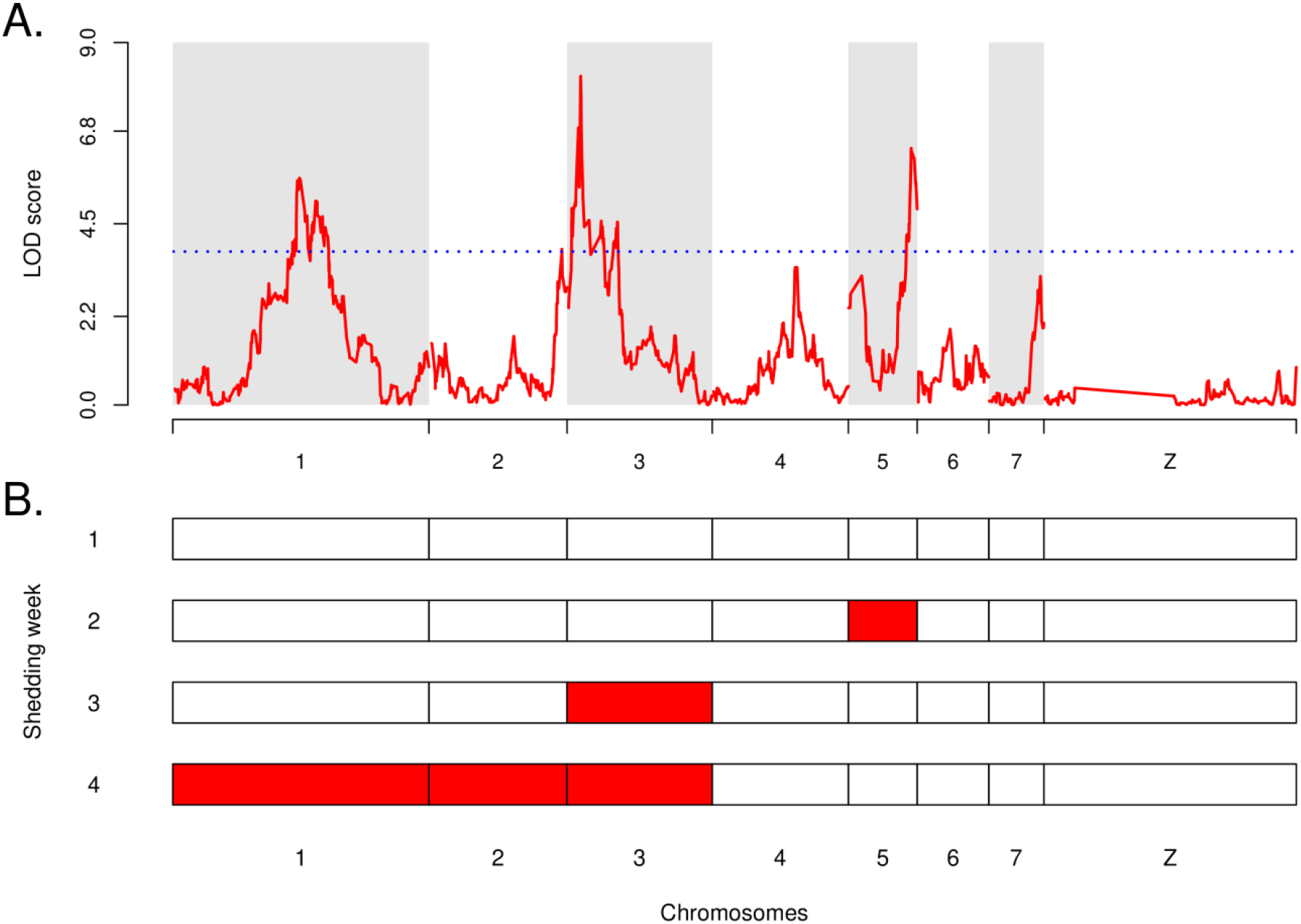
Linkage analysis of the transmission stage production in *S. mansoni* parasites and QTLs interactions. **(A)** Linkage analysis between the average cercarial production phenotype and F2 *S. mansoni* genotypes (combined crosses A and B) demonstrated that transmission stage production is a polygenic trait controlled by three major QTLs with statistically significant LOD: on chr. 1 (LOD= 5.63), on chr. 2 (LOD= 8.16) and on chr. 5 (LOD= 6.37). The blue dotted line represents the 1,000 permutation threshold. **(B)** Genetic architecture of cercarial production is influenced by the shedding week (first shedding week corresponds to week 4 post-infection), demonstrating a sequential pattern of QTL emergence involved in transmission stage production. On shedding week 1, none of the QTLs identified pass the permutation threshold, while on shedding week 2, cercarial production is linked to QTL on chr. 5. On shedding week 3, the main QTL linked to cercarial production is on chr. 3. Finally, on shedding week 4, the QTL on chr.1 predominates, along with QTLs on chr. 2 and chr. 3.

We used a two-dimensional genome scan to search for additional contributing QTLs and to identify interactions among loci. We found two minor QTLs on chromosome 2 (LOD= 3.87) and 4 (LOD= 3.32) which contributed significantly to explain the phenotype variance. The two-QTL scan demonstrated that all five QTLs act additively: we found no evidence for epistasis between the loci (Table 1). Together, these 5 QTLs explained 28.56% of the variation in cercarial production, with a full LOD score of 29.79 (Table 2). The three major QTLs explained most of the phenotype: chr. 1 = 8.10% (*p* = 5.49 × 10^™10^), chr. 3 = 6.10% (*p* = 8.49 × 10^™8^), chr. 5 = 5.57% (*p* = 3.31 × 10^™7^). The two minor QTLs explained less phenotypic variation: chr. 2 = 3.80% (*p*= 3.38 × 10^™5^) and chr. 4 = 3.52% (*p*= 7.06 × 10^™5^) but these contributions are highly significant.

**Table 1:** Summary table of the two-dimensional, two-QTL genome scan identifying QTL interactions (additive or epistatic effects) between the QTL involved in cercarial production in *Schistosoma mansoni* parasite. LOD values are statistically significant when they are above the 5% threshold LOD for epistatic (interaction) and/or additive effect. We have demonstrated here additive interaction of various loci involved in transmission stage production but no epistatic interaction.

| Pairs of loci | Position locus 1 | Position locus 2 | LOD interaction (epistatis) | LOD additive |
| --- | --- | --- | --- | --- |
| **5% threshold** | - | - | **5.68** | **5.84** |
| Chr1:Chr2 | 43.84 | 45.94 | 0.611 | **10.48** |
| Chr1:Chr3 | 43.84 | 5.44 | 0.186 | **12.40** |
| Chr1:Chr4 | 43.84 | 28.95 | 1.020 | **9.05** |
| Chr1:Chr5 | 43.84 | 21.56 | 1.367 | **11.88** |
| Chr2:Chr4 | 45.94 | 28.90 | 0.738 | **7.48** |
| Chr2:Chr5 | 45.94 | 21.56 | 1.244 | **9.60** |
| Chr3:Chr5 | 5.44 | 21.56 | 2.587 | **10.96** |
| Chr4:Chr5 | 28.90 | 21.56 | 0.622 | **9.20** |

**Table 2:** QTLs interactions modeling (additive model) and estimation of the proportion of the cercarial production variance (%var) explained by each of the loci involved in the phenotype. All together, the 5 QTLs explained 28.56% of variation in the transmission stage production in *S. mansoni* parasites.

| Loci | LOD | %var | P value |
| --- | --- | --- | --- |
| **Full additive model (all 5 QTLs)** | **29.8** | **28.56** | **0** |
| Chr1 | 9.52 | 8.10 | **5.49x10<sup>-10</sup>** |
| Chr2 | 4.6 | 3.80 | **3.38x10<sup>-5</sup>** |
| Chr3 | 7.26 | 6.10 | **8.49x10<sup>-8</sup>** |
| Chr4 | 4.26 | 3.52 | **7.06x10<sup>-5</sup>** |
| Chr5 | 6.66 | 5.57 | **3.31x10<sup>-7</sup>** |

### Genetic control of shedding varies across time

When we conducted the linkage analysis using the cercarial quantities produced each week rather than the average across all four weeks (i.e. from week 4 to week 7 post-exposure, see Figure 1), we observed a sequential pattern of QTL emergence (Figure 3B, Supplementary figure 2). On shedding week 1, none of the identified QTLs passed the permutation threshold (Figure 3B, Supplementary figure 2A), while on shedding week 2, the QTL on chr. 5 predominated and a QTL on chr. 3 had arisen (Figure 3B, Supplementary figure 2B). On shedding week 3, the main QTL was on chr. 3 (Figure 3B, Supplementary figure 2C). Finally, on shedding week 4, the QTL on chr. 1 predominated, along with QTLs on chr. 2 and 3 (Figure 3B, Supplementary figure 2D). Hence, the parasite genes that determine cercarial production vary across time during the snail infection.

### Allelic inheritance and impact on cercarial production in S. mansoni *parasites*

For each QTL, we determined the interactions between alleles from L and H shedding parents by analyzing the average production of cercariae per genotype (i.e. LL: homozygous for the low shedding allele, LH: heterozygous and HH: homozygous for the high shedding allele; Figure 4). Our analysis showed codominance for the chr. 2, 3 and 5 QTLs: F2 progeny with LH genotype produced more cercariae than LL genotype but less HH genotype (Figure 4). However, for the chr. 1 and 4 QTLs showed patterns consistent with recessive inheritance. At these loci, F2 progeny with HH genotype produced more cercariae (pairwise comparison using Wilcoxon rank sum test; chr. 1: HH vs. LH: *p* = 4.2 × 10^™5^ and HH vs. LL: *p* = 8.2 × 10^™6^; chr. 4: HH vs. LH:, *p* = 0.0109 and HH vs. LL: *p* = 0.0034) than the other genotypes (i.e. LL and LH; pairwise comparison using Wilcoxon rank sum test; chr. 1: LL vs. LH, *p* = 0.22; chr. 4: LL vs. LH, *p* = 0.56; Figure 4).

**Figure 4.**
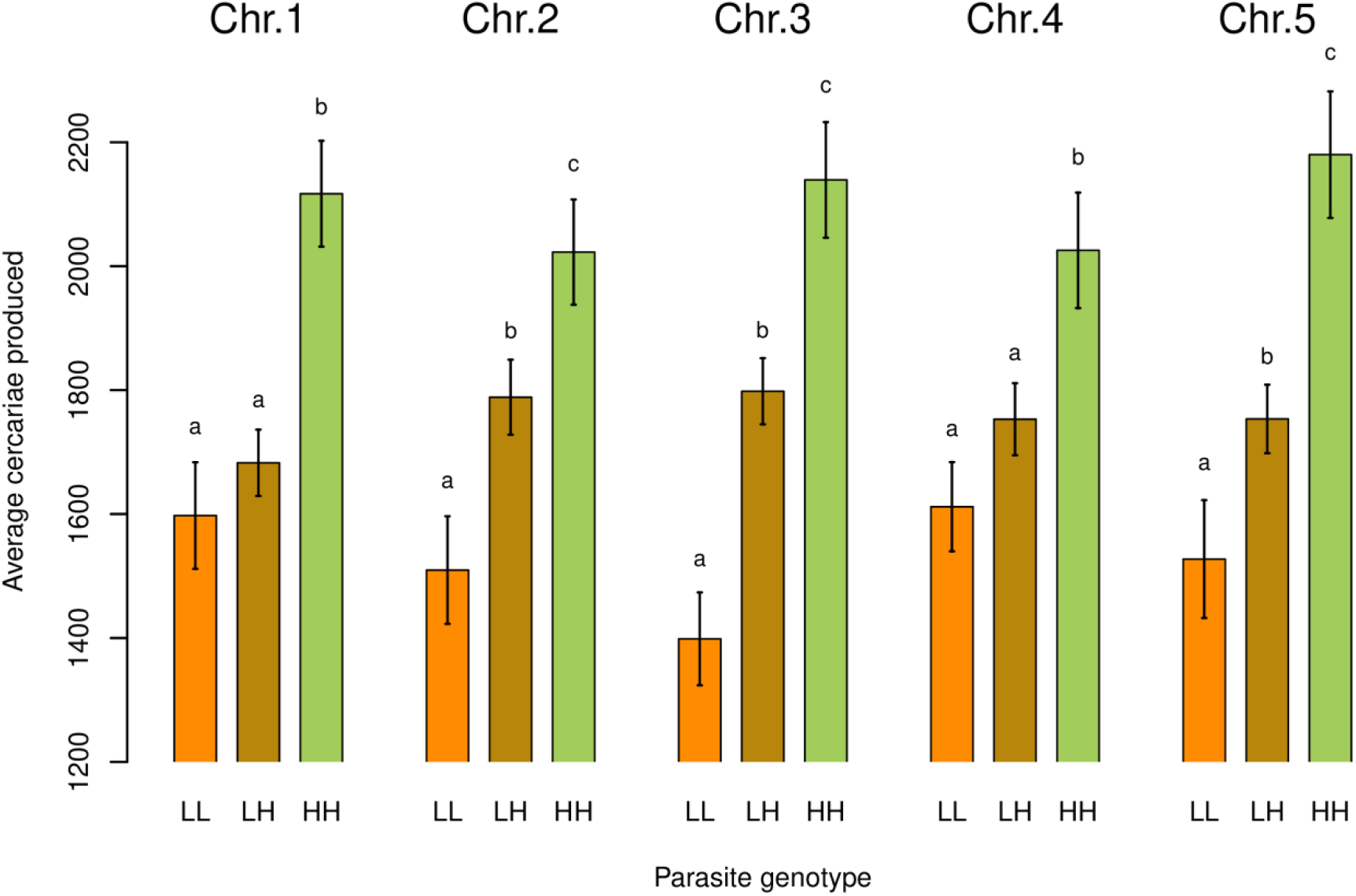
Inheritance of transmission stage production alleles. Impact of the parasite genotype on the average number of cercariae produced for each major and minor QTL linked to transmission stage production. On chr. 1, cercarial production is not significantly different when parasites are homozygous for the “low shedding” allele (i.e. LL) or heterozygous (i.e. LH) but only when parasites are homozygous for the “high shedding” allele (i.e. HH). High shedding allele (H) is recessive and low shedding allele (L) is dominant. On chr. 2, cercarial production is significantly different for all the three parasite genotypes encountered (i.e. LL, LH and HH). High and Low shedding alleles act co-dominantly. On chr. 3, we observed the same results as exhibited for chr. 2. For this locus, High and Low shedding alleles also act co-dominantly. For the locus located on chr. 4, we encounter the same scenario as for the locus on chr. 1: the cercarial production is not significantly different for parasites exhibited LL or LH genotypes but for the HH genotypes. H allele is recessive and L dominant. Finally, on chr. 5, we observed similar results as already shown for chr. 2 and 3, where the cercarial production is significantly different whatever the parasite genotype. L and H alleles are co-dominant for this locus. Parasite genotypes (i.e. LL, LH, HH) not connected by the same letter for a given locus are significantly different (post-hoc test).

### Candidate genes controlling transmission stage production in S. mansoni *parasite*

We listed the genes located within the 1.8 LOD-support interval of each QTL and expressed either in daughter sporocysts or in cercariae. We identified 314 genes on chr. 1, 87 on chr. 2, 31 on chr. 3, 76 on chr. 4 and 86 on chr. 5 (Table 3 and Supplementary Table 1). We then prioritized genes using an objective index (“genoscore”, see Material & Methods section), that accounts for the genotypes of the parents (fix alternative alleles), the presence of non-synonymous mutations and their impact on the protein predicted by snpEff (see Material & Methods section), the CDS length and the distance from the QTL peak. Using this objective approach, we identified the three most likely candidate genes within each QTL that may be involved in transmission stage production (Table 4).

**Table 3:** Summary table of the total number of genes located within the 1.8 LOD-support interval of each QTL peak, and expressed into the different parasite stages: daughter sporocysts only, cercariae only, both stages or not expressed at all.

| Loci | Total number of genes | Genes expressed in daughter sporocysts only | Genes expressed in cercariae only | Genes expressed in both stages | Genes not expressed |
| --- | --- | --- | --- | --- | --- |
| **Chr1** | 329 | 297 | 301 | 284 | 15 |
| **Chr2** | 94 | 87 | 83 | 78 | 2 |
| **Chr3** | 32 | 28 | 28 | 25 | 1 |
| **Chr4** | 79 | 70 | 72 | 66 | 3 |
| **Chr5** | 94 | 78 | 82 | 74 | 8 |

**Table 4:** Summary table of the 3 best candidate genes for each locus involved in cercarial production in the parasite *S. mansoni* based on the computed genoscore CDS (see Materials & Methods) and their relative positions to the maximum peak. NA: gene not captured by the *S. mansoni* baits (exome capture). In bold: selected best candidate genes based on their annotations (GFF or HHsearch).

| Chromosome | Gene ID | GFF annotation | HHsearch annotation | Max. LOD | Genoscore |
| --- | --- | --- | --- | --- | --- |
| Chr1 | Smp_152850 | Tetratricopeptide-like helical domain superfamily | **Putative 70 kda peptidylprolyl isomerase PFL2275c** | 5.63 | 1.646 |
|  | Smp_057230 | Aminoacyl-tRNA synthetase, class II | Seryl-tRNA synthetase (SerRS) | NA | 1.286 |
|  | Smp_246820 | *No annotation available* | hypothetical protein TTHA1254 | 4.05 | 1.114 |
| Chr2 | Smp_347740 | Coiled-coil domain-containing protein 39 | **Nucleoporin_FG2** | 3.86 | 7.657 |
|  | Smp_328700 | *No annotation available* | Engineered transmembrane domain variant | NA | 2.153 |
|  | Smp_170430 | FAD dependent oxidoreductase | D-aminoacid oxidase, N-terminal domain | 2.81 | 1.491 |
| Chr3 | Smp_343200 | Pleckstrin homology domain | **G-protein coupled receptor kinase 2 (beta-adrenergic receptor kinase 1)** | 8.16 | 1.653 |
|  | Smp_190770 | Lanthionine synthetase C-like | Nisin biosynthesis protein NisC | 5.41 | 1.426 |
|  | Smp_138060 | *No annotation available* | Cationic trypsin (e.c.3.4.21.4) | 7.66 | 1.227 |
| Chr4 | Smp_313330 | *No annotation available* | *No annotation available* | 3.35 | 1.732 |
|  | Smp_150840 | **Zinc finger, C2H2, LYAR-type** | 60S ribosomal protein L8, 60S | 2.47 | 1.502 |
|  | Smp_334250 | Epithelial sodium channel | Acid-sensing ion channel 1 | 3.42 | 1.257 |
| Chr5 | Smp_330680 | *No annotation available* | *No annotation available* | NA | 6.997 |
|  | Smp_330700 | *No annotation available* | *No annotation available* | NA | 3.784 |
|  | Smp_330760 | **IQ motif, EF-hand binding site** | **Calmodulin, IQ domain-containing protein G** | 4.50 | 3.631 |

The genes prioritization on each of the three major QTLs revealed three compelling candidates: (i) peptidylprolyl isomerase (chr. 1) which regulates many biological processes, including intracellular signaling, transcription, inflammation, immunomodulation and apoptosis (Ramachandran et al., 2014; Wei et al., 2013), (ii) G-protein coupled receptor kinase 2 (chr. 3) which is involved in cell migration (Randazzo et al., 2000) and (iii) a protein with unknown function (chr. 5) which suggests schistosome specific factor yet to be characterized. On chr. 5, another candidate, the Calmodulin IQ domain protein, is of interest: this protein is a calcium sensor and can stimulate changes in the actin cytoskeleton mediated by proteins such as myosin (Houdusse and Cohen, 1996).

In the two minor QTLs, the top candidate genes encode a nucleoporin (chr. 2) and a protein of unknown function (chr. 4). The second best candidate gene on chr. 4 is also of interest: it encodes a LYAR, a cell growth regulating nucleolar protein.

### Linkage analysis of snail physiological trait associated with transmission stage production

Laccase-like activity and hemoglobin rate in the *Biomphalaria* snail hemolymph were correlated (Supplementary figure 3C and 3F). These metrics provide proxies to evaluate snail health and the impact of schistosome infection (Le Clec’h et al., 2016; Le Clecʼh et al., 2019). Both parameters were negatively correlated with F2 cercarial production (laccase-like activity: Pearson’s correlation test, *p* = 4.31 × 10^™15^, correlation coefficient = −0.56; Hemoglobin rate: Pearson’s correlation test, *p* = 8.014 × 10^™12^, correlation coefficient = −0.50; Supplementary figure 3D and 3E). There was no correlation with the F1 cercariae production (laccase-like activity: Pearson’s correlation test, *p* = 0.426, correlation coefficient = −0.09; Hemoglobin rate: Pearson’s correlation test, *p* = 0.661, correlation coefficient = - 0.05; Supplementary figure 3A and 3B). This was expected because F1s were heterozygous for markers from each parent at all five QTLs and show limited variation in cercarial production.

We performed a linkage analysis to investigate parasite genes linked to laccase-like activity and the hemoglobin rate in the snail host hemolymph. We did not find significant QTLs across *S. mansoni* genome linked to these snail hemolymph phenotypes.

## DISCUSSION

### How many loci are involved in transmission/virulence?

We determined the genetic architecture of transmission stage production trait in schistosome parasites using classical linkage mapping. We showed that transmission/virulence traits in schistosome are polygenic, involving three major and 2 more minor loci. These results are consistent with the idea that relatively small numbers of genes are involved in strongly selected traits such as virulence/transmission. The obvious caveat here is that loci of small effect may be difficult to identify even in well powered linkage analyses.

Three other studies have used linkage mapping to dissect the genetic basis of parasite transmission/virulence trait (Table 5). In the protozoan parasite *Toxoplasma gondii*, four sets of genetic crosses were used to identify the genetic basis of acute virulence between different clonal lineages from North America (Behnke et al., 2014). These crosses identified several QTLs linked to acute virulence and resulted in the identification of a set of polymorphic genes in the GTPase serine/threonine kinase family in secretory organelles called rhoptries (Behnke et al., 2016). In another protozoan model, the rodent malaria parasite (*Plasmodium yoelii yoelii*), Pattaradilokrat *et al*. (2009) and Otsuki *et al*. (2009) showed that just one locus on chr. 13 was linked to parasite growth rate and host virulence. Plant parasite nematode system provides a directly comparable macroparasite system in which virulence has been examined using linkage mapping. In the root-knot nematode *Meloidogyne hapla*, crosses between parasites that differ in ability to produce galls on the roots of the ornamental nightshade *Solanum bulbocastanum* identified three QTLs linked to nematode transmission success (Thomas and Williamson, 2013).

**Table 5:** Do transmission/virulence traits tend to be oligogenic (i.e. controlled by few genes)? Summary table of the number of loci and genes involved in transmission/virulence or growth life-history traits for various eukaryotic parasite systems (reviewed from the literature).

| Parasite system | Phenotypes | Number of loci involved (Chrom. #) | Number of gene(s) involved (Gene name) | Percentage of phenotypic variations explained by the loci | References |
| --- | --- | --- | --- | --- | --- |
| *Toxoplasma gondii* | Acute virulence | 1 (XII) | 1 (ROP5) | - | Behnke et al., 2011 |
|  | Acute virulence | 3 (VIIa/VIIb/XII) | 3 (ROP18/ROP16/ROP5) | - | Reese et al., 2011; Saeij et al., 2006 |
|  | Acute virulence | 2 (VIIa/Ia) | 1 (ROP18) | - | Fentress et al., 2010; Taylor et al., 2006 |
|  | Growth | 4 (VIIa/XI/XII/Ia) | - | - | Yamamoto et al., 2011 |
|  | Migration | 1 (VIIa) | - | - | Taylor et al., 2006 |
|  | Acute virulence | 1 (XII) | 1 (ROP5) | - | Behnke et al., 2015 |
| *Plasmodium yoelii yoelii* (rodent malaria parasite) | Growth rate | 1 (XIII) | 1 (*pyebl*) | - | Otsuki et al., 2009; Pattaradilokrat et al., 2009 |
| *Meloidogyne hapla* (plant pathogen nematode) | Egg mass number | 2 (VII/XIII) | - | - | Thomas and Williamson, 2013 |
|  | Total eggs among F2 lines | 2 (IV/VII) | - | - | Thomas and Williamson, 2013 |
| *Schistosoma mansoni* | Transmission stage production/Virulence | 5 (3 major: I/III/V and 2 minor: II/IV) | 5 potential best candidate genes (peptidylprolyl isomerase/ G-protein coupled receptor kinase / Calmodulin and Nucleoporin_FG2/ LYAR cell growth regulating nucleolar protein) | 28.56% | *Present study* |

In all these three cases, relatively few loci control transmission/virulence traits, as we see in schistosome parasites (Table 5). This question of “few loci or genes” versus “many loci or genes” controlling key life-history traits is fundamental to our understanding of adaptation in nature. As demonstrated by an increasing number of studies, few large-effect loci (and subsequently genes) have been shown to drive rapid adaptation (reviewed in Messer et al., 2016).

### How do parasite genes impact numbers of cercariae shed?

Genes may impact numbers of cercariae shed from infected snails by (i) influencing growth and differentiation of sporocysts, (ii) changing the balance of investment between cercarial production and daughter sporocyst production, (iii) influencing the ability of cercariae to escape from the snail host (Figure 5). Two lines of evidence strongly suggest that the genes determining cercarial shedding may act on at least different processes.

**Figure 5.**
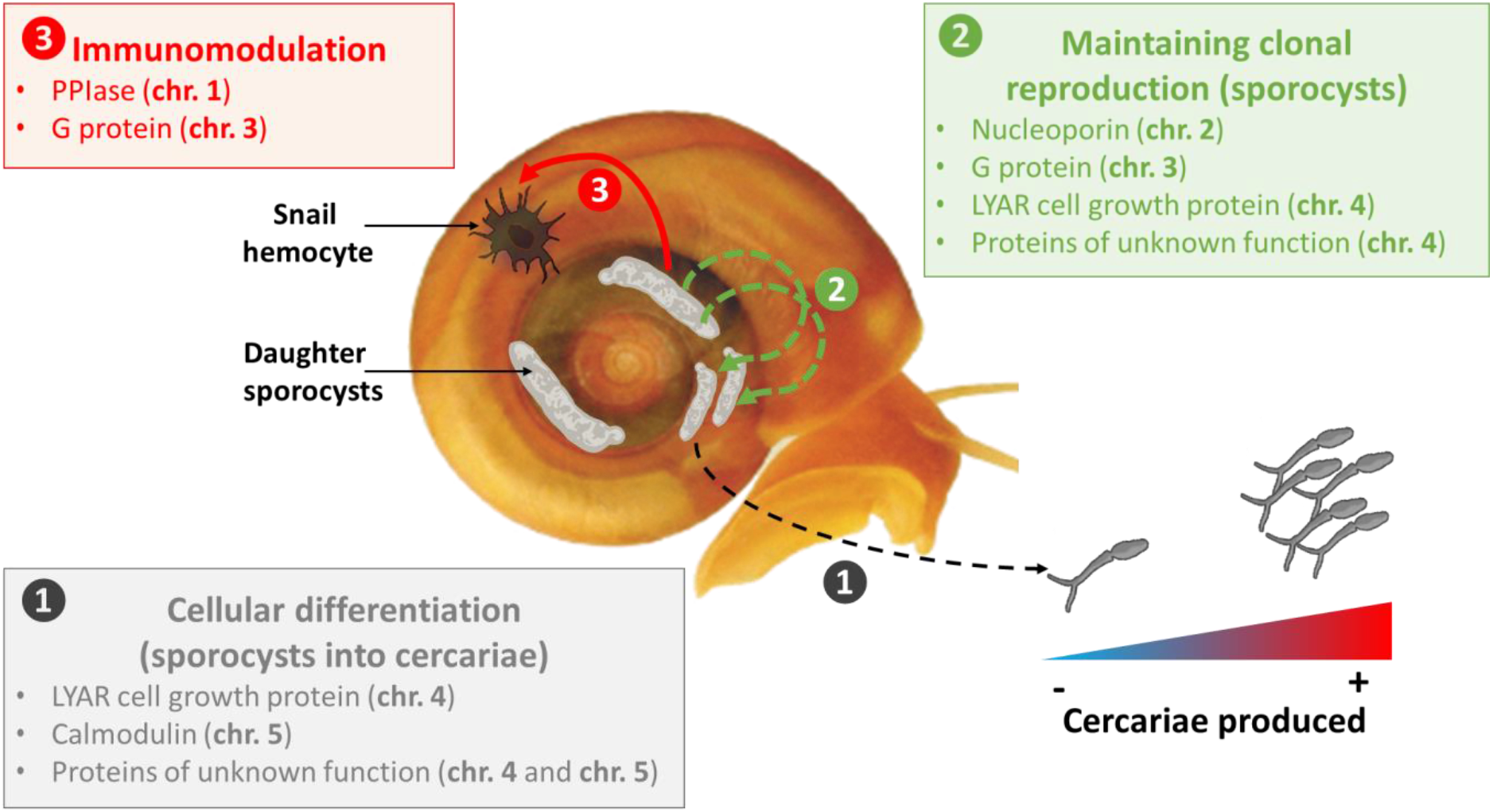
Potential mechanisms involved in production of schistosome transmission stage. Our genetic analysis of transmission stage production and virulence in *S. mansoni* parasites revealed that this complex trait is controlled by 5 loci with potentially different and complementary mechanisms involved to induce and maintain cercarial production over time. This could involve (i) first the differentiation of sporocysts cells into cercariae through the Calmodulin calcium sensor (chr. 5) (ii) then the maintenance of the clonal reproduction of sporocysts and the continuous cercarial production through the G protein (chr. 3) and the Nucleoporin (chr. 2), and finally (iii) the modulation of the snail immune response to protect parasitic cells through the PPIase (chr. 1) and the G protein (chr. 3). The LYAR cell growth protein (chr. 4) could also be involved all along the patent period to regulate the growth and multiplication of sporocysts cells and the development of cercariae. The proteins of unknown function are also of high interest: they could reveal new mechanisms involved in transmission/virulence and will deserve further attention.

First, we previously compared sporocyst size and growth kinetics in the parental populations using qPCR. SmLE-H sporocysts comprise on average 47% of cells within infected snails, while SmBRE-L sporocysts comprise 25% of cells within infected snails (Le Clecʼh et al., 2019). However, this is not sufficient to fully explain the difference in cercariae produced by these two populations, because the SmBRE-L infected snails shed significantly fewer cercariae than predicted from qPCR measures of sporocysts cells in infected snails. These results are consistent with independent action of genes to determine sporocyst size and cercarial production.

Second, when we dissected the genetic architecture of cercarial production over the 4-week patent period of the infection, we observed sequential emergence of QTLs. This is also consistent with the idea that genes influencing cercarial shedding may act on different processes to determine this cercarial production. This sequential emergence of loci could also explain the evolution of sporocysts kinetics trajectories observed in parental population (Le Clecʼh et al., 2019). Moreover, we highlighted that the high shedding alleles acted co-dominantly with the low shedding alleles at QTLs on chr. 2, chr. 3 and chr. 5 but were recessive at QTLs on chr. 1 and chr. 4. These differences in allelic inheritance could also support the existence of different mechanisms involved in the modulation of transmission stage production and virulence: co-dominant loci would involve alleles with dose effect while recessive loci would involve loss-of-function alleles. In the next section we evaluate potential candidate genes and speculate at which stage of the parasite these may play a role.

### Candidate genes underlying transmission/virulence traits

The first QTL linked to cercarial production appears on the second shedding week and is located on chr. 5. The three leading candidate genes under this QTL include two genes encoding proteins of unknown function and a gene encoding a calmodulin (CAM) IQ domain protein. The proteins of unknown function are of high interest as they could reveal new mechanisms involved in transmission/virulence and will deserve further attention. CAM IQ domain proteins belong to calcium sensors and can stimulate changes in the actin cytoskeleton mediated by proteins such as myosin (Houdusse and Cohen, 1996). In schistosome, the calcium sensor (CAM IQ domain protein) could modulate the differentiation of daughter sporocysts cells into cercariae and induce the cercarial shedding early in the patent period. Calcium binding proteins are known to play an important role in amoebic pathogenesis and are essential for the *Entamoeba histolytica* parasite growth (Sahoo et al., 2004), while in *Toxoplasma gondii* parasite, CAM-like proteins are essential and contribute to regulate parasite motility and host cell invasion (Long et al., 2017).

During the third shedding week, the major QTL is on chr. 3, where the leading candidate gene is a G-protein coupled receptor kinase 2. In *Entamoeba histolytica* G-protein are involved in pathogenesis-related cellular processes, such as migration, invasion, phagocytosis and evasion of the host immune response by surface receptor capping (Bosch and Siderovski, 2013). In *S. mansoni*, this G-protein coupled receptor kinase 2 may stimulate sporocyst expansion or renewal into snail host tissues (i.e. new generation of daughter sporocysts) while escaping host immune defenses, to maintain the parasitic infection and eventually, cercarial production.

During the fourth shedding week, QTLs on chr. 1, chr. 2 and chr. 3 are involved in transmission stage production. The leading candidate gene on chr. 1 is a peptidylprolyl isomerase (PPIase). Most PPIases characterized in parasites belong to the cyclophilin family (Ünal and Steinert, 2014) and are strong immunomodulatory proteins (Wang and Heitman, 2005). In the apicomplexan *Theileria*, a secreted prolyl isomerase modulates host leukocyte transformation (Marsolier et al., 2015); similar immunomodulatory peptidylprolyl isomerases have been demonstrated in two other apicomplexan parasites, *Toxoplasma gondii* and *Neospora caninum* (Tuo et al., 2005). PPIases have been shown to be primary actors of host-parasite interactions and are certainly essential for the development and differentiation of parasitic protozoans (e.g *Leishmania* parasites), which show a high degree of plasticity in their cellular organization and metabolic status during their infection cycles (Yau et al., 2010). In our schistosome model, PPIase may modulate snail host immune response to infection, while maintaining proliferation and viability of sporocysts cells. The best candidate gene under the chr. 2 QTL is a nucleoporin, protein that is essential to microtubule organization and dynamics during mitosis. In the parasite *Plasmodium falciparum*, nucleoporins are essential for parasite proliferation in human erythrocytes (Dahan-Pasternak et al., 2013). Nucleoporins are also essential in the transport of mRNA from the nucleus to the cytoplasm after transcription and are involved in cell migration (Chatel and Fahrenkrog, 2012). In schistosome, these proteins might trigger the sporocysts clonal proliferation and cercariae production.

The QTL on chr. 4, revealed by the two-dimensional genome scan, did not pass the permutation threshold at any of the 4-week patent period of the infection. Therefore, this QTL could have a minor but continuous effect throughout the patent period. Interestingly, the leading candidate genes under this QTL encode protein of unknown function followed by a LYAR cell growth regulating nucleolar protein. Proteins of unknown function are again of high interest and will deserve further attention. In addition, the LYAR protein, required for cell proliferation (Hui Li et al., 2009) and highly present in early embryos of mammals (Su et al., 1993), could also be involved all along the patent period to regulate the growth and multiplication of schistosome sporocysts cells and the development of cercariae.

### Prospects for functional analysis

Figure 5 summarizes potential involvement of candidate genes in cercarial shedding in *S. mansoni*. Functional validation is required to investigate involvement of these candidate genes. While RNAi methods are available for gene knockdown in adult worms (Krautz-Peterson et al., 2010), delivery of RNA to the sporocysts within the snail is challenging. We have tried several approaches to induce RNAi in daughter sporocysts including injection of *ds* or *si*RNA into the snail hemolymph/tissues, but have so far not been able to knockdown genes in this stage. CRISPR provide a possible alternative approach but is still at an early stage of development for *S. mansoni* (Ittiprasert et al., 2019; Sankaranarayanan et al., 2020; You et al., 2021). Once effective gene manipulation approaches are available for late sporocysts we will be able to directly examine the involvement of our leading candidate genes.

Stem cells play a central role in schistosome development and reproduction. Wang and colleagues (2018) have demonstrated that primary sporocysts containing different cellular types including stem cells. In our model, we suspect that SmLE-H and SmBRE-L sporocysts may exhibit different cellular trajectories, with differences in development of cell that differentiate to generate cercariae and those that give birth to the next generation of daughter sporocysts (Collins, 3rd et al., 2011). Advances in our understanding of stem-cell differentiation of *S. mansoni* within the molluscan host coupled with single-cell transcriptomics of daughter sporocysts from our two populations provides and alternative approach to investigate the molecular basis of these transmission-related developmental differences at the cellular and molecular levels (Collins, 3rd et al., 2011; Wang et al., 2018, 2013). Functional and cell biology characterization of this key life history trait underlying transmission and virulence will be the focus of future work on this system.

## MATERIALS & METHODS

### Ethics statement

This study was performed in accordance with the Guide for the Care and Use of Laboratory Animals of the National Institutes of Health. The protocol was approved by the Institutional Animal Care and Use Committee of Texas Biomedical Research Institute (permit number: 1419-MA).

### Biomphalaria glabrata s*nails and* Schistosoma mansoni *parasites*

Uninfected inbred albino *Biomphalaria glabrata* snails (line Bg26 derived from 13-16-R1 line; Bonner et al., 2012) were reared in 10-gallon aquaria containing aerated freshwater at 26-28°C on a 12L-12D photocycle and fed *ad libitum* on green leaf lettuce. All snails used in this study had a shell diameter between 8 and 10 mm, as snail size can influence cercarial production (Gérard C., Moné H., 1993; Tavalire et al., 2016). For all the experiments presented in this study, we used inbred snails to minimize the impact of snail host genetic background on the parasite life history traits (Le Clecʼh et al., 2019).

The SmLE-H schistosome (*Schistosoma mansoni*) population (high shedder, H) was originally obtained from an infected patient in Belo Horizonte (Minas Gerais, Brazil) in 1965 and has since been maintained in laboratory (Lewis et al., 1986), using *B. glabrata* NMRI and Bg26 population as intermediate host and Syrian golden hamster (*Mesocricetus auratus*) as definitive hosts. The SmBRE schistosome population (low shedder, L) was sampled in 1975 from Recife (East Brazil) (Theron et al., 2014) and has been maintained in the laboratory in its sympatric albino Brazilian snail host (BgBRE) using hamsters or mice as the definitive host.

### Genetic crosses between high shedder and low shedder S. mansoni *parasites*

Schistosomes are sex separated parasites that simplifies crosses (Valentim et al., 2013). In order to identify QTL(s) controlling the number of cercariae produced by schistosome parasites, we conducted crosses between parasites from our high shedder (SmLE-H) and low shedder (SmBRE-L) populations (Le Clecʼh et al., 2019). We performed two reciprocal genetic crosses (cross A: male SmBRE (L) x female SmLE (H); cross B: female SmBRE (L) x male SmLE (H)) to i) replicate our cross experiment and ii) to test for the potential influence of sex chromosomes on the transmission phenotype (Figure 1).

Design of the crosses is summarized in Figure 1. To obtain the parental (F0) parasite generation we exposed individual snails (192 per parasite populations) to single miracidia in 24 well-plates overnight. Exposed snails were then maintained in trays (48 per tray) for 4 weeks as described in Le Clec’h *et al*. (2019). At four weeks post-exposure, we placed each snail in a well of a 24 well-plate, added in 1 mL freshwater to each well and placed the plate under artificial light for 2 h to induce cercarial shedding. Shedding was conducted late morning to early afternoon every week from week 4 to week 7 post-infection. To track cercarial production of individual snails over the 4 weeks of the patent period, we isolated each infected snail in a uniquely labeled 100 mL glass beaker filled with ∼50 mL freshwater at the first shedding, fed them *ad libitum* with fresh lettuce and kept them in the dark during the 4 weeks (except when shedding was induced). We determine parasite gender by PCR (Chevalier et al., 2016) on cercariae collected from week 4, for the two schistosome populations (H and L). One female and one male were randomly selected for producing the next generation. At week 5, cercarial shedding was induced and 2 female golden Syrian hamsters per cross were exposed to 250 cercariae of the female genotype and 250 cercariae of the male genotype. For cross A, the male genotype was SmBRE-L and the female genotype was SmLE-H. For cross B, the male genotype was SmLE-H and the female genotype was SmBRE-L. After 45 days, we euthanized and perfused the hamsters to recover the F0 adult male and female worms for DNA extraction and sequencing. We also collected the livers containing the eggs.

We applied the same procedures for the F1 generation. We hatched F1 miracidia from eggs recovered from hamster livers following Le Clec’h *et al*. (2019). For each cross, we exposed individual Bg26 snails (288 per cross) to single F1 miracidia. At week 4, we isolated infected snails in individual beakers for tracking individual shedding. We collected cercariae of the first shedding for parasite gender determination. At week 5, we infected 4 female hamsters per cross with 250 cercariae of the females and the male genotypes of the selected F1s. We euthanized and perfused hamsters 45 days post infection to recover F1 adult worms and collected livers containing the F2 eggs.

After hatching F2 miracidia from eggs recovered from hamster livers, we exposed 2,000 individual Bg26 snails (1,000 per cross) to single miracidia. After 4 weeks, we isolated the infected snails in individual beakers as above to track individual shedding. We collected all the cercariae of a snail from the first or the second shedding in individual microtubes for further DNA extractions.

### Life history traits measured on S. mansoni parasites and virulence toward its snail hosts

We measured larval output (i.e. cercarial production) of F0, F1 and F2 parasites. We used the same inbred snail population for all infections to minimize the impact of host snail genetic variation (Figure 1). All snails were infected with single miracidia to allow examination of cercarial shedding from single parasite genotypes. We also measured the impact of these parasitic infections on the snail host by quantifying snail survival and physiological responses (i.e. laccase-like activity and hemoglobin rate in the hemolymph) at end point (Figure 1). The phenotypic datasets are available on Zenodo (DOI: 10.5281/zenodo.4383248).

#### a. S. mansoni cercarial production over time

We placed infected snails in 1 mL freshwater in a 24 well plate under artificial light (as described above) every week for 4 weeks (week 4 to 7). After replacing the snail back to its beaker, we sampled three 10 (for the high shedder parasites) or 100 µL (for the low shedder parasites) aliquots of each well, added 20 µL of 20X normal saline and counted the immobilized cercariae under a microscope. We multiplied the mean of the triplicated measurement by the dilution factor (10) to determine the number of cercariae produced by each infected snail. We assessed cercarial production every week from week 4 to 7 post-exposure in snails infected with F0, F1 and F2 parasites.

#### b. Snail physiological response to parasitic infection

During week 7, 3 days after the last cercarial shedding, we collected hemolymph from all surviving snails infected with F0 parental populations (i.e. SmLE-H and SmBRE-L), F1s and F2s (Le Clec’h et al., 2016). We measured both laccase-like activity and the hemoglobin (protein carrying oxygen in *B. glabrata* hemolymph) rates in the hemolymph of each infected snail (Le Clecʼh et al., 2019).

### Estimation of the minimum number of loci influencing transmission stage production

Castle and Wright proposed an estimate *n*_*e*_ of the minimum effective number of genetic factors explaining trait segregation in crosses between to two lines based on the phenotypic mean and variance (Castle, 1921; Wright and Morton, 1968).

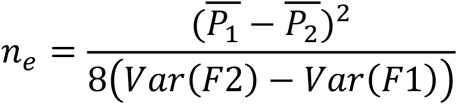

where *P* is the phenotypic mean of the parents (i.e. *P*_1_ for SmLE-H and *P*_2_ for SmBRE-L), *Var(F2)* is the phenotypic variance of the F2 population progeny and *Var(F1)* is the phenotypic variance of the F1 population. We calculated *n*_*e*_ for each cross independently.

### Whole genome and exome sequencing of schistosome parasite

We sequenced genomes from F0 parent worms, and exomes from F1 worms used to generate the F2 and from 188 F2s cercariae for both cross A and cross B.

#### a. gDNA extraction

We extracted gDNA from F0 and F1 worms and from F2 cercariae using the Blood and Tissue kit (Qiagen), following the manufacturer protocol, with minor modifications. We homogenized worms in DNA extraction kit lysis buffer using sterile micro pestles. For F2 cercariae, frozen samples were thaw at 4°C, centrifuged 5 minutes at x300 *g* to pellet the cercariae. We removed the water supernatant, added lysis buffer to the pellet, and vortexed to homogenize. We incubated at 56°C worm and cercariae samples for 2 and 1 hour, respectively. We recovered gDNA in 200 µL of elution buffer. We quantified the worm samples gDNA using the Qubit dsDNA HS Assay Kit (Invitrogen) while F2 cercarial DNA was directly whole genome amplified to provide sufficient DNA for exome capture.

#### b. Whole genome amplification (WGA) of F2 cercarial samples

We performed WGA on each F2 gDNA cercarial sample using the Illustra GenomiPhi V2 DNA Amplification kit (GE Healthcare Life Sciences, USA) according to Le Clec’h *et al*. (2018). We quantified the WGA DNA using the Qubit dsDNA HS Assay Kit (Invitrogen).

#### c. Whole genome and exome library preparation and sequencing

We prepared whole genome libraries of F0s using the KAPA HyperPlus kit (KAPA Biosystem) according to the manufacturer’s protocol. For each F0 library, we sheared 500 ng of gDNA by adaptive focused acoustics (Duty factor: 10%; Peak Incident Power: 175; Cycles per Burst: 200; Duration: 180 seconds) in AFA tubes, using a Covaris S220 instrument with SonoLab software version 7 (Covaris, Inc., USA), to recover fragmented DNA between 150-200 bp. We used 6 PCR cycles for post-ligation library amplification.

We captured F1 and F2 *S. mansoni* exomes using the SureSelect^XT2^ Target Enrichment System (Agilent). The design of the custom baits used to capture the *S. mansoni* exome (SureSelect design ID: S0398493) is described in Chevalier *et al*. (2014) and exome capture methodology follows Le Clec’h *et al*. (2018).

We sequenced the libraries on a HiSeq 2500 sequencer (Illumina) using 100 bp pair-end reads. On each sequencing lane, we either pooled 32 exome capture libraries or 2 whole genome libraries. Raw sequence data are available at the NCBI Sequence Read Archive under accession numbers PRJNA667697 (reviewer link: https://dataview.ncbi.nlm.nih.gov/object/PRJNA667697?reviewer=bo7f89g1actpms73hb12r2n8).

### Bioinformatic analysis

Jupyter notebook and scripts used for processing the sequencing data and mapping QTLs are available on Github (https://github.com/fdchevalier/Cercarial-production).

#### a. Sequence analysis and variant calling

We aligned the sequencing data against the *S. mansoni* reference genome (schistosoma_mansoni.PRJEA36577.WBPS14) using BWA (v0.7.17) (Li and Durbin, 2009) and SAMtools (v1.10) (Heng Li et al., 2009). We used GATK (v4.1.8) (DePristo et al., 2011; McKenna et al., 2010) to mark PCR duplicates and recalibrate base scores. We used the HaplotypeCaller module of GATK to call variants (SNP/indel) and the GenotypeGVCFs module to perform a joint genotyping on each chromosome or unassambled scaffolds. We merged VCF files using the MergeVcfs module. We annotated the variants using snpEff (v.4.3.1t) (Cingolani et al., 2012). All these steps were automatized using snakemake (v5.14.0) (Köster and Rahmann, 2018).

#### b. In silico schistosome sexing

Sexual determination in schistosome relies on Z and W chromosomes: females are ZW while males are ZZ. We can already determine sex molecularly by PCR using sex specific markers (Chevalier et al., 2016). We decided to take advantage of the sequencing data to perform *in silico* sexing by comparing the read depth along the Z chromosome. The Z chromosome carries a Z-linked region (located between 11 and 44 Mb) which never recombines with the W chromosome contrariwise to the rest of the Z chromosome (pseudo-autosomal region). While the Z-linked region in males (ZZ) will display the same read depth as the pseudo-autosomal region, the Z-linked region in females (ZW) will display only half of the read depth of the pseudo-autosomal region (Supplementary Figure 4). We can therefore determine a read depth ratio between the Z-linked and pseudo-autosomal regions: a ratio around 1 will correspond to a male carrying two Z chromosomes while a ratio around 0.5 will correspond to a female carrying only one Z chromosome. We computed the read depth at each base of the chromosome Z using SAMtools and BEDtools (v2.29.0) (Quinlan and Hall, 2010). We used R to first smooth the read depth data which may show local high variation using the runmed function on 101 contiguous sites, then we kept only site showing a read depth of more than 5 to finally compute the ratio on Z-linked over Z pseudo-autosomal read depth. We validated this approach by comparing molecular and *in silico* sexing of some F2s.

#### c. Linkage analysis

We first reduced the VCF file using the SelectVariants module of GATK to include sites called in at least half of the samples. The QTL analysis was performed with R (v3.5.1) (R Core Team, 2018). We loaded the VCF file using vcfR (v1.10.0) (Knaus and Grünwald, 2017), filtered out sites with genotype quality (GQ) less than 30, a read depth (DP) less than 10 and with more than 20% of missing data. We retained alternatively fixed variants between parents (i.e., fully informative variants) and converted the genotype data into an R/qtl compatible format. We then run the scanone function of R/qtl (v1.46.2) (Broman et al., 2003) to map QTL in each cross or in the combined crosses using the expectation– maximization (EM) algorithm. We performed 1,000 permutations for each test.

Having identified QTLs, we selected a subset of around 400 markers total (number of markers per chromosome being proportional to the chromosome size) from the combined crosses and tested QTL interaction using the scantwo function from the same package with 1,000 permutations. We used a subset of markers because this function is computationally intensive. We tested the allelic dominance on the combined crossed using a custom R code.

#### d. Candidate gene prioritization

To identify candidate genes involved in cercarial production, we first reduced the VCF file to sites within QTL regions identified previously. We loaded the VCF file in R using vcfR and refined the QTL regions by determining the chromosomal positions corresponding to a drop of 1.8 LOD from the highest LOD score of each QTL. We then retained only genotypes from the cross parents and filtered out sites with a GQ lower than 30 and a DP lower than 6. We used a custom score (genoscore) to objectively prioritize candidate genes in each QTL region:

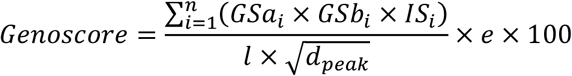

The genoscore is computed either by gene or gene’s coding sequence (CDS). For each site i of a gene or CDS, we first computed a genotype score (GS) for each cross a and b by associating a score of 20 if genotypes were alternatively fixed in each cross parent or a score of 5 if fixed in one parent and unknown in the other (any other genotype combination received 0) and an impact score (IS) from the snpEff annotation (low impact or modifier mutation = 1, moderate impact = 5, high impact = 10). We summed the results for the *n* sites present in the gene or CDS. We weighed this sum using the length *l* of the gene or the CDS, and the square root of the distance *d* between the position of the highest LOD score (peak) and the start of the gene or the CDS. This was finally modulated by an expression factor *e* of 1 if the gene was expressed in mature sporocysts from shedding snails (Buddenborg et al., 2019) or in cercariae (Protasio et al., 2012) or −1 otherwise. Jupyter notebook and scripts used to obtain gene expression are available on Github (https://github.com/fdchevalier/Sm_gene_expression).

#### e. Gene annotations

To complete the gene annotation of the GFF file, we ran the HHsearch tool from the HH-suite (v3.3.0-0) (Steinegger et al., 2019) on each predicted protein sequences. This method relies on generating hidden Markov models (HMM) for a given sequence and compares HMM-HMM alignments. We compared our sequences to three databases (pdb70, scop70 and pfam) and selected the best match if the probability generated by HHsearch was at least of 50%. Gene identifications from the GFF file and the HHsearch analysis were combined in a table and used during the analysis of gene candidates. Jupyter notebook and scripts used for this analysis are available on Github (https://github.com/fdchevalier/Sm-GFF-HHPred-table).

### Statistical analysis

All statistical analyzes and graphs were performed using R software (v3.5.1) (R Core Team, 2018). When data were not normally distributed (Shapiro test, p < 0.05), we compared results with non-parametric Kruskal-Wallis test followed by pairwise Wilcoxon-Mann-Whitney post-hoc test or a simple pairwise comparison Wilcoxon-Mann-Whitney test. When data followed a normal distribution, we used one–way ANOVA or a pairwise comparison Welsh *t*-test. We performed correlation analysis using Pearson’s correlation test. The confidence interval of significance was set to 95% and *p*-values less than 0.05 were considered significant.

## ACKNOWLEDGEMENTS

We thank Michael S. Blouin (Oregon State University) for the Bg26 snail line, Guillaume Mitta and Benjamin Gourbal (University of Perpignan France) for SmBRE *S. mansoni* population and Philip LoVerde (UT Health San Antonio) for the SmLE *S. mansoni* population. We thank the Vivarium of the Southwest National Primate Reasearch Center (SNPRC) for providing rodent care. This research was supported by a Cowles fellowship (WL) from Texas Biomedical Research Institute (13-1328.021), and NIH R01AI133749 (TJCA) and was conducted in facilities constructed with support from Research Facilities Improvement Program grant C06 RR013556 from the National Center for Research Resources. SNPRC research at Texas Biomedical Research Institute is supported by grant P51 OD011133 from the Office of Research Infrastructure Programs, NIH.

WL, FDC and TJCA designed the experiments. WL, FDC, MMW, VM and GAA performed the experiments. WL and FDC performed the data analyses. WL and TJCA drafted the manuscript. All authors read and approved the final manuscript.

## COMPETING INTERESTS

The authors declare that they have no competing interests.

## SUPPLEMENTARY FIGURES

**Supplementary figure 1:**
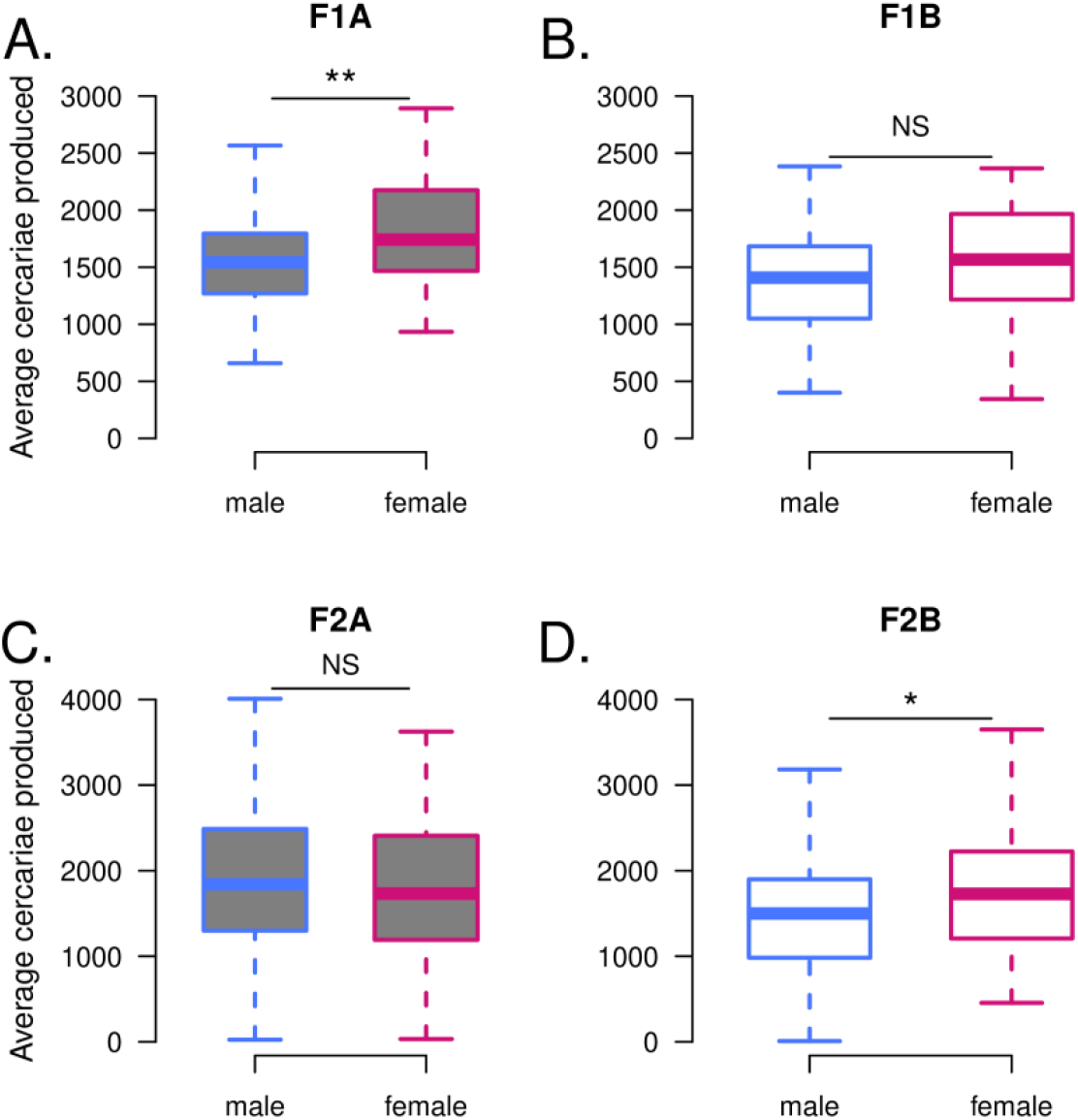
Impact of *S. mansoni* gender on the cercarial production for F1s (A-B) and F2s (C-D). **(A)** Male sporocysts produced significantly less cercariae than female sporocysts in F1A cross parasite but **(B)** there is no difference driven by the gender of the parasites for the F1B cross. For F2 progeny, gender has no impact on cercarial production in cross A **(C)**, but male sporocysts produced significantly less cercariae than female sporocyts in cross B **(D)**. NS: No significant difference in cercarial production between the two considered groups; \**p* < 0.05; ** *p* ≤ 0.02; *** *p* ≤ 0.002.

**Supplementary Figure 2.**
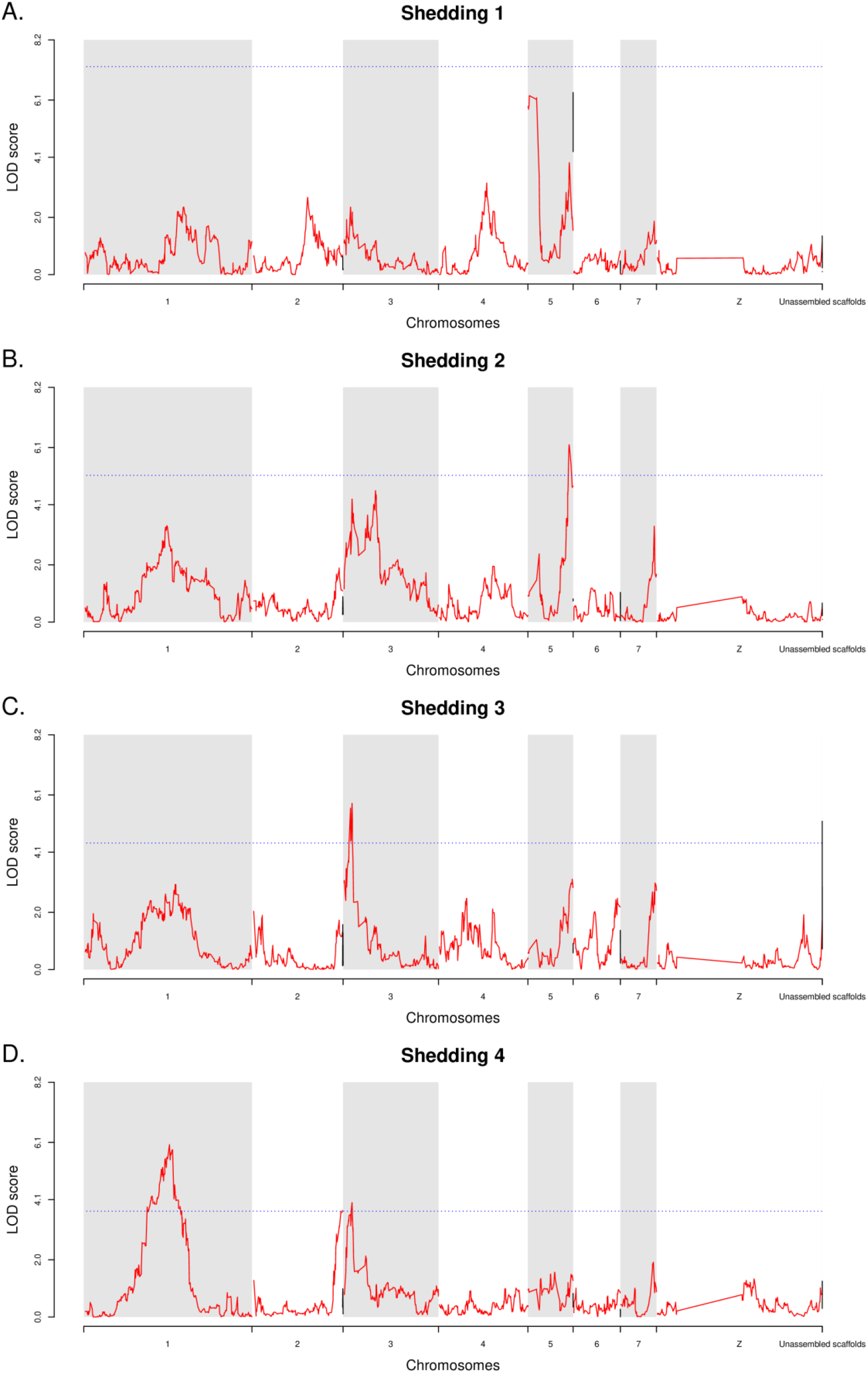
Influence of the shedding week on the genetic architecture of cercarial production. **(A)** On shedding week 1, none of the QTLs identified pass the permutation threshold **(B)** On shedding week 2, the only QTL involved in the transmission trait is located on chr. 5. **(C)** On shedding week 3, QTL on chr. 3 is the only one with a significant LOD score. **(D)** Finally on shedding week 4, QTL on chr. 1 seems to be the one explaining most of the phenotype, along with QTL on chr. 2 and on chr. 3. Red line refers to assembled chromosome and black line to unassembled scaffolds assigned to chromosome or still unassigned. The blue dotted line corresponds to the 1,000 permutation threshold.

**Supplementary Figure 3.**
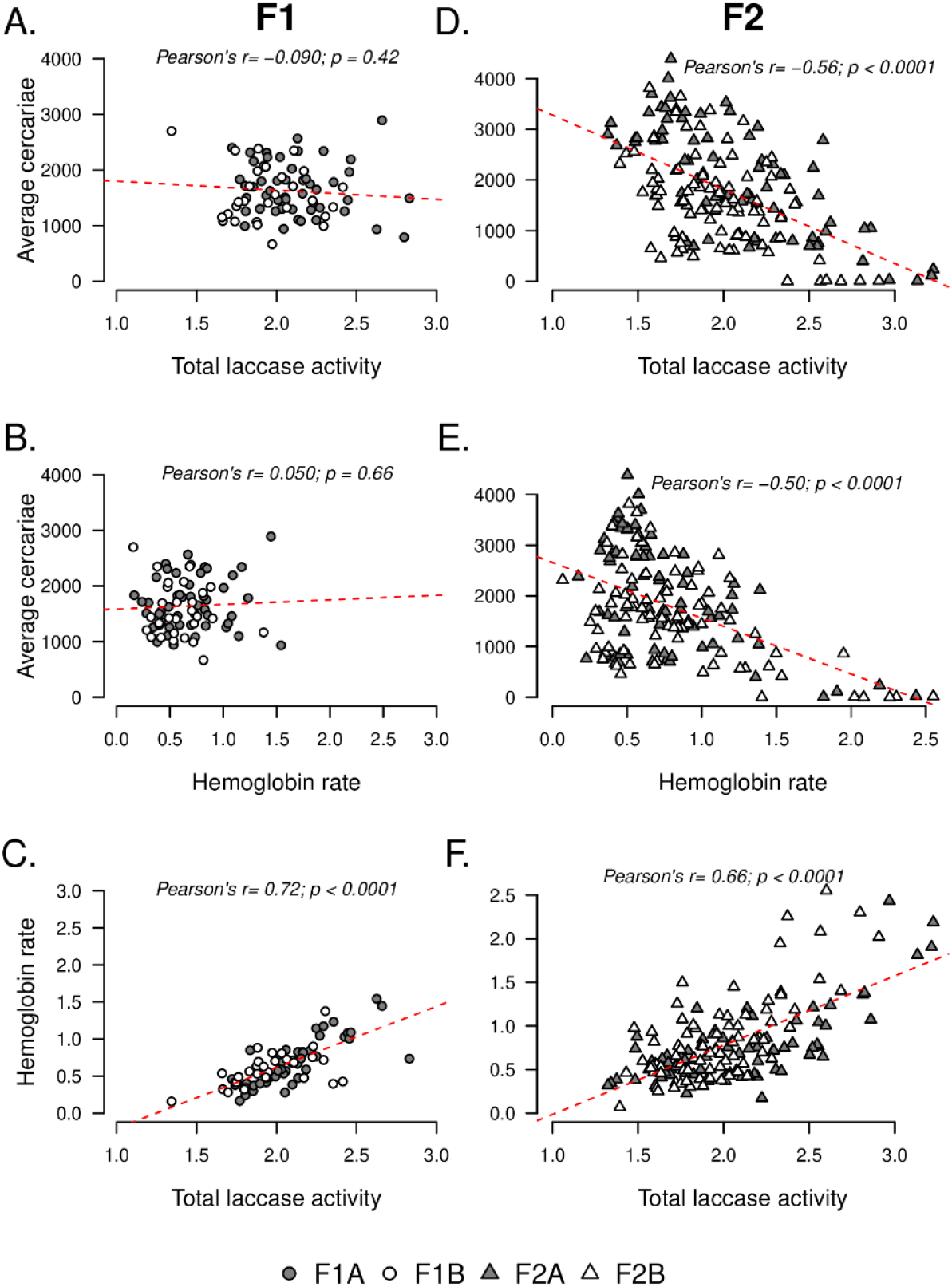
Correlation between laccase-like activity and hemoglobin rate measured in the hemolymph of infected intermediate snail host *Biomphalaria glabrata* and the cercarial production for F1 (A-C) and F2 (D-F) *S. mansoni* parasite progeny (7.5 weeks post-infection). We did not observe significant correlations neither **(A)** between laccase-like activity and the average production of cercariae, nor **(B)** between the hemoglobin rate and the average cercarial production in F1 *S. mansoni* progeny. This result was expected as all the individual exhibited heterozygous genotype (i.e. LH) and showed an intermediate phenotype in their transmission stage production compared to their respective parents. However, we observed a significant negative correlation **(D)** between the average cercariae produced by F2 parasite progeny and the total laccase-like activity in the hemolymph of the infected snail (Pearson’s test, coef. = −0.56, *p* < 0.0001) and **(E)** between these cercarial production and the hemoglobin rate measured in infected snail hemolymph (Pearson’s test, coef. = −0.50, *p* < 0.0001). For both snail physiological parameters measured in the hemolymph, we highlighted a strong positive correlation (For F1: Pearson’s test, coef. = 0.72, *p* < 0.0001 and for F2: Pearson’s test, coef. = 0.66, *p* < 0.0001) whatever the parasite generation analyzed.

**Supplementary Figure 4.**
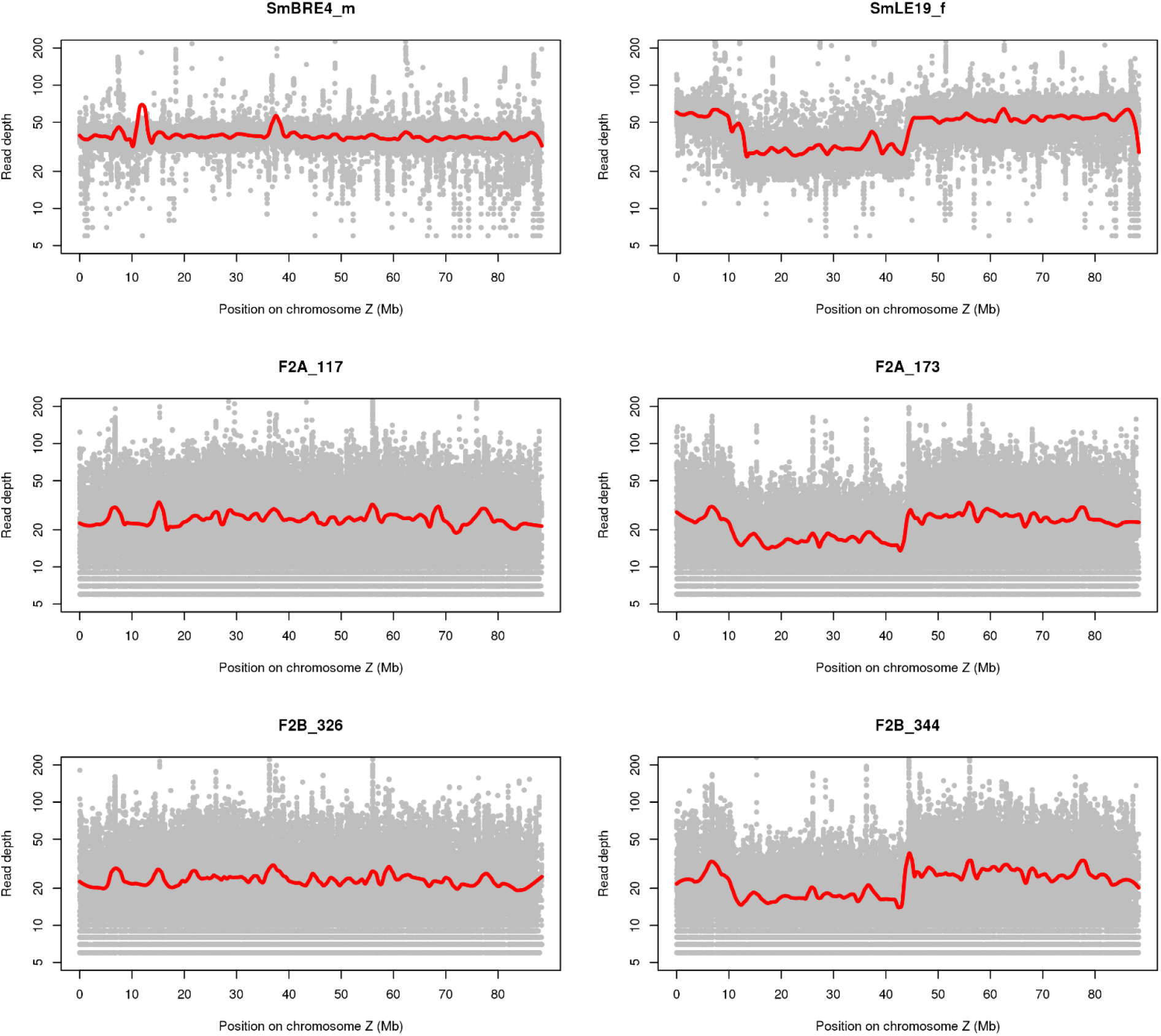
Example of read depth across the Z chromosome in males and females. The left column represents male samples and the right column female samples. The top rows are known sex samples (parents of cross A), the two other rows are samples for which we predicted sex *in silico*. The grey dots represent the smoothed read depth, the red line the local regression of the read depth obtained from the loess function.

**Supplementary Table 1. Complete list of candidate genes linked to transmission stage production in *S. mansoni* parasite.** This table gives summary information for each gene under the QTL regions and detailed information about the variants in these regions.

## Notes

https://zenodo.org/record/4383248

### Competing Interest Statement

The authors have declared no competing interest.

